# Methylene Blue-Photodynamic Therapy Effectively Kills Antibiotic-Resistant Bacteria from Pediatric Patients with Perforated Appendicitis

**DOI:** 10.1101/2025.04.29.651230

**Authors:** Korry T. Wirth, Eric Y. Wang, Darrian S. Hawryluk, Matthew M. Byrne, Marjorie J. Arca, Derek S. Wakeman, Walter Pegoli, David G. Darcy, Martin S. Pavelka, Nicole A. Wilson, Timothy M. Baran

**Affiliations:** Division of Pediatric Surgery, Department of Surgery, University of Rochester Medical Center; Department of Microbiology and Immunology, University of Rochester; Department of Biomedical Engineering, University of Rochester; Department of Imaging Sciences, University of Rochester Medical Center

## Abstract

**Objectives:** Perforated appendicitis (PA) is the most common cause of intra-abdominal abscess in children and is associated with higher costs, extended hospitalizations, and worse postoperative outcomes compared to uncomplicated appendicitis. Photodynamic therapy (PDT) utilizes a photosensitizer and light to generate cytotoxic reactive species that are efficacious against bacteria. We sought to determine whether PDT could effectively eliminate bacteria isolated from the peritoneal cavities of pediatric patients with perforated appendicitis.

**Hypothesis:** In monoculture planktonic samples, PDT achieves a minimum of 99.9% bacterial cell kill (3 log_10_ growth reduction per mL).

**Methods:** After IRB approval, bacterial subcultures from 30 pediatric surgical patients with PA were developed as planktonic monocultures. Samples were derived from standard clinical collection of peritoneal fluid in patients undergoing surgery for PA. Cultures were incubated with the photosensitizer methylene blue (MB) and exposed to a 665 nm laser to perform PDT. Control conditions were MB alone, laser alone, or no MB and no laser. Log reductions in bacterial growth between groups were compared using two-way ANOVA.

**Results:** Bacteria were isolated from 28 (93.3%) specimens, and 26 (86.7%) were polymicrobial. The most prevalent organisms included *Escherichia coli* (76.7%), the *Streptococcus anginosus* group (56.7%), *Bacteroides fragilis* (46.6%), and *Pseudomonas aeruginosa* (26.7%), with antibiotic resistance common among E. *coli* isolates.

For *E. coli*, MB-PDT caused a 5.86 ± 0.39 log_10_ reduction compared to the no-treatment control (p<0.001). For *S. anginosus* and *P. aeruginosa*, MB-PDT resulted in reductions of 5.91 ± 0.88 log_10_ and 2.23 ± 0.64 log_10_, respectively (all p<0.001). PDT showed comparable efficacy in isolates with multiple antibiotic resistance relative to those that were antibiotic pan-susceptible for *E. coli* (p=0.760), *S. anginosus* group (p>0.999), and *P. aeruginosa* (p=0.991).

**Conclusions:** PDT application to bacterial isolates from patients with perforated appendicitis achieved >99.9% bacterial kill in all isolated strains other than *P. aeruginosa*, regardless of antibiotic susceptibility, suggesting that PDT may be a viable adjunct to *in vivo* antimicrobial therapy.

## Introduction

Appendicitis is the most common pediatric surgical emergency^1,2^. Appendicitis begins as acute inflammation secondary to blockage of the appendiceal orifice, which leads to congestion, necrosis, and eventual perforation of the appendiceal wall. Unfortunately, 25-30% of pediatric cases in the United States present with perforated appendicitis (PA)^3,4^. Compared with non-perforated appendicitis, PA is associated with higher costs, more extended hospitalizations, and worse postoperative outcomes^3,5–8^. The prolonged length of stay of patients with PA primarily is typically due to the time required for intravenous antibiotics to control the intra-abdominal infection caused by stool and bacteria leakage from the appendiceal perforation. This extended hospitalization places a significant burden on the healthcare system. The reported mean hospital length of stay for PA ranges from 5 to 15 days^9,10^ and amounts to median total hospital costs of $17,000-$22,500 per patient^11^. Patient- and family-level strains may include physical, mental, and social burdens as well as financial impact of loss of income and out of pocket expenses. Alternative treatment strategies for PA are therefore needed to mitigate the direct and indirect costs of complications due to infection.

One such potential treatment is photodynamic therapy (PDT), which uses photosensitive drugs to generate reactive oxygen species upon exposure to sub-thermal levels of visible light. PDT is effective against various microbial species^12^, including antibiotic-resistant strains^13,14^, and does not result in acquired resistance^15^. PDT is a localized therapy that requires the presence of both photosensitizer and treatment light to cause oxidative damage. This is in contrast to systemic antibiotic treatment, which may affect the host microbiome, potentiate antimicrobial resistance, and may result in adverse medical events^16–18^, which is of particular concern in pediatric patients, where early antibiotic exposure is frequent and has been linked to both adverse health outcomes^19–21^ and the proliferation of antimicrobial resistance genes^22,23^.

Particular success with PDT has been shown using the photosensitizer methylene blue (MB), which is FDA-approved for treating methemoglobinemia^24^. MB-PDT has been used clinically to treat superficial infections of the skin and oral cavity^25–28^, and has also been applied preoperatively to reduce surgical site infections in adult spine surgery^29^. Thus, MB use is deemed safe in humans.

We have previously demonstrated the efficacy of MB-PDT in treating a limited subset of bacteria common in abdominal infections^30,31^. However, these prior studies did not include bacteria isolated from patients with perforated appendicitis. Since bacterial properties can vary based on their source, clinical setting and genetic differences, evaluating PDT efficacy against bacteria isolated directly from the site or pathophysiologic process of eventual clinical application is essential for translation to clinical practice. For example, we have shown that in *Staphylococcus aureus*, mutations related to oxidant response can affect PDT efficacy^32^.

The current study aimed to characterize the bacteria present in intra-abdominal fluid aspirated during appendectomy from pediatric patients with perforated appendicitis and to evaluate the efficacy of MB-PDT in treating these bacteria. To achieve this, bacteria isolated from aspirated fluid were cultured planktonically and treated with MB-PDT at fixed drug and light doses. This was followed by the evaluation of growth *in vitro* 24 hours post-PDT. We hypothesized that MB-PDT would result in at least 99.9% bacterial kill (3 log_10_ reduction relative to control), with potential variability in efficacy across isolated species. The results of this study are the first to investigate MB-PDT as a novel therapy that could change the current surgical paradigm for perforated appendicitis and intra-abdominal abscesses.

## Materials and Methods

### Patient population

The Research Subjects Review Board at the University of Rochester Medical Center approved all study procedures. Prior to sample collection, informed consent was obtained from the parent or legal guardian of all participants, and informed assent was obtained from all patients greater than 8 years of age.

Samples were collected from 30 pediatric patients undergoing routine appendectomy for perforated appendicitis. Inclusion criteria were: (1) preoperative diagnosis of suspected perforated appendicitis, (2) age less than 18 years, and (3) parent or legal guardian able to understand the study and provide written consent. Exclusion criteria were: (1) absence of appendiceal perforation at the time of operation, (2) insufficient quantity of peritoneal fluid for collection, and (3) known infectious diseases such as HIV, hepatitis C, or tuberculosis.

### Sample collection and processing

Study data were obtained from biospecimens collected as standard of care during routine appendectomy for PA. Peritoneal fluid was aspirated and collected in a sterile container, a step that is standard of care for appendectomy for perforated appendicitis at our institution.

Peritoneal fluid was sent directly to the Clinical Microbiology laboratory at the University of Rochester Medical Center for routine processing and evaluation, which included bacterial identification and antibiotic susceptibility testing. Subject-de-identified bacterial cultures were sub-cultured by Clinical Microbiology and delivered to the study team for *in vitro* PDT experiments. Samples were cataloged and stored at −80 ºC for subsequent experiments.

### Photodynamic therapy conditions

For *in vitro* experiments, individual bacterial subcultures were streaked out onto BHI agar plates (Criterion Brain Heart Infusion (BHI) Broth with 1.5% BD Bacto Agar) and incubated overnight at 37°C. *S. capitis* and *S. anginosus* group were streaked onto blood plates (Difco Columbia Broth with 1.5% BD Bacto Agar and 5% HemoStat Laboratories Defib Sheep Blood) and incubated in candle jars overnight to promote more robust growth. One day before the experiment, 25 mL BHI liquid cultures were inoculated using these plates and incubated overnight at 37°C.

On the day of the experiment, liquid cultures were harvested via centrifugation (5000x g,15 minutes at 4°C). Pellets were washed twice in 25 mL phosphate-buffered saline (PBS) using the same settings. Samples were then adjusted to an optical density of 1 at 600 nm (OD_600_). 1 mL aliquots were transferred to microcentrifuge tubes. 300 μg/mL MB (Akorn, Inc., Lake Forest, IL) was added to samples in the MB groups (MB+) and subsequently incubated in a dark room on a rotator for 30 minutes. For control samples without MB (MB−), MB was replaced with an equivalent volume of PBS and incubated under the same conditions. All samples then underwent two rounds of centrifugation (10,000 x g, 10 minutes) and PBS washing. Bacterial samples were then transferred into twelve-well plates. One was sent for illumination (L+), while the other was shielded from treatment light (L−).

Treatment light was delivered by a lens-tipped optical fiber connected to a 665 nm laser (LDX-3230-665, LDX Optronics, Inc., Maryville, TN), with output power set to achieve a fluence rate of 4 mW/cm^2^ to a total fluence of 7.2 J/cm^2^; resulting in a total illumination time of 30 minutes. We have previously shown efficacy against multiple bacterial species using this combination of MB concentration and light parameters^30,31^.

In total, each isolated bacterial strain was treated in four groups. The experimental group was treated with MB and light (MB+L+). In addition, we also performed control experiments under the following conditions (Figure 1): no MB, light only (MB−L+); MB only, no light (MB+L−); and no light, no MB (MB−L−). For MB-free controls (MB−L+ and MB−L−), MB was replaced with an equivalent volume of PBS, and light-free controls (MB+L− and MB−L−) were shielded from treatment light.

**Figure 1.**
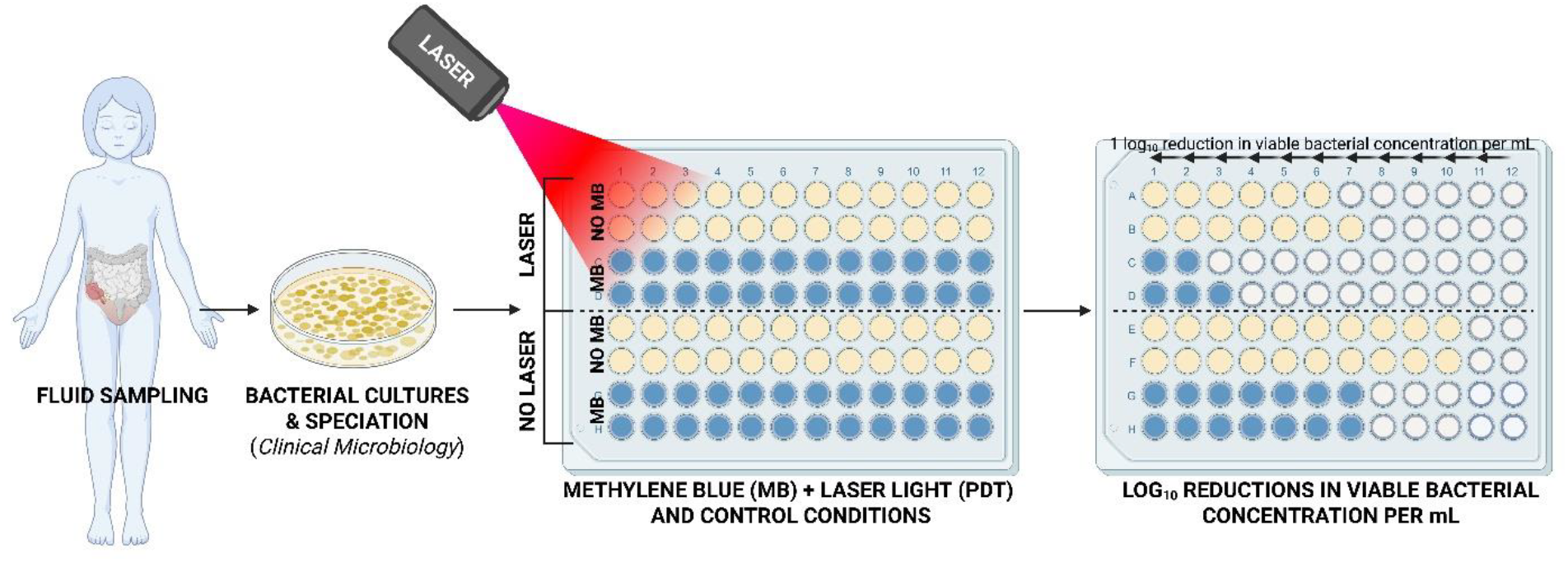
Schematic of Experimental Protocol.

After treatment under PDT or control conditions, 20 μL of each sample was transferred to a 96-well plate with 180 μL of BHI in each well. Samples were serially diluted across the plate and incubated at 37 ^°^C overnight. Subsequently, the plates were analyzed for growth, with each consecutive well corresponding to a 1 log_10_ increase in bacterial growth (Figure 1). Therefore, results are reported as differences in log_10_ viable bacterial concentration per mL.

### Statistical analysis

For each bacterial isolate, differences in bacterial burden were compared between the no treatment (MB−L−), control conditions (MB+L− and MB−L+), and MB-PDT (MB+L+) conditions using two-way ANOVA, with Tukey’s post hoc test for multiple comparisons, unless indicated otherwise. Summary values are presented as mean ± standard deviation. All reductions are reported with respect to the no-treatment condition unless otherwise specified. Statistical analysis was performed in GraphPad Prism (v10.2.1, GraphPad Software, LLC, Boston, MA).

## Results

A total of 85 bacterial isolates were obtained from 30 subjects. The mean subject age was 8.4 ± 3.8 years, and 73.3% (22/30) of subjects were male. 66.7% (20/30) of subjects identified as White, 6.7% (2/30) as Black or African American, 6.7% (2/30) as Asian, 10% (4/30) as multiracial, and 0% did not report race. 6.7% (2/30) were of Hispanic or Latino ethnicity.

### Bacteria isolated

Bacteria were isolated in 93.3% (28) of specimens. Of the 30 subjects, 86.7% (26) had a polymicrobial infection, 6.7% (2) had a monomicrobial infection, and 6.7% (2) showed no bacterial growth from the aspirated peritoneal fluid sample. The most prevalent bacteria (Figure 2A) were *Escherichia coli* (73.3%, 22/30), *Streptococcus anginosus group* (60.0%, 18/30), *Bacteroides fragilis* (46.6%, 14/30), and *Pseudomonas aeruginosa* (26.7%, 8/30). Of the 22 patients with *E. coli*, 27 unique *E. coli* strains were isolated. There were 18 unique *S. anginosus* strains isolated from 18 patients, and 9 unique *P. aeruginosa* isolated from 8 patients.

**Figure 2.**
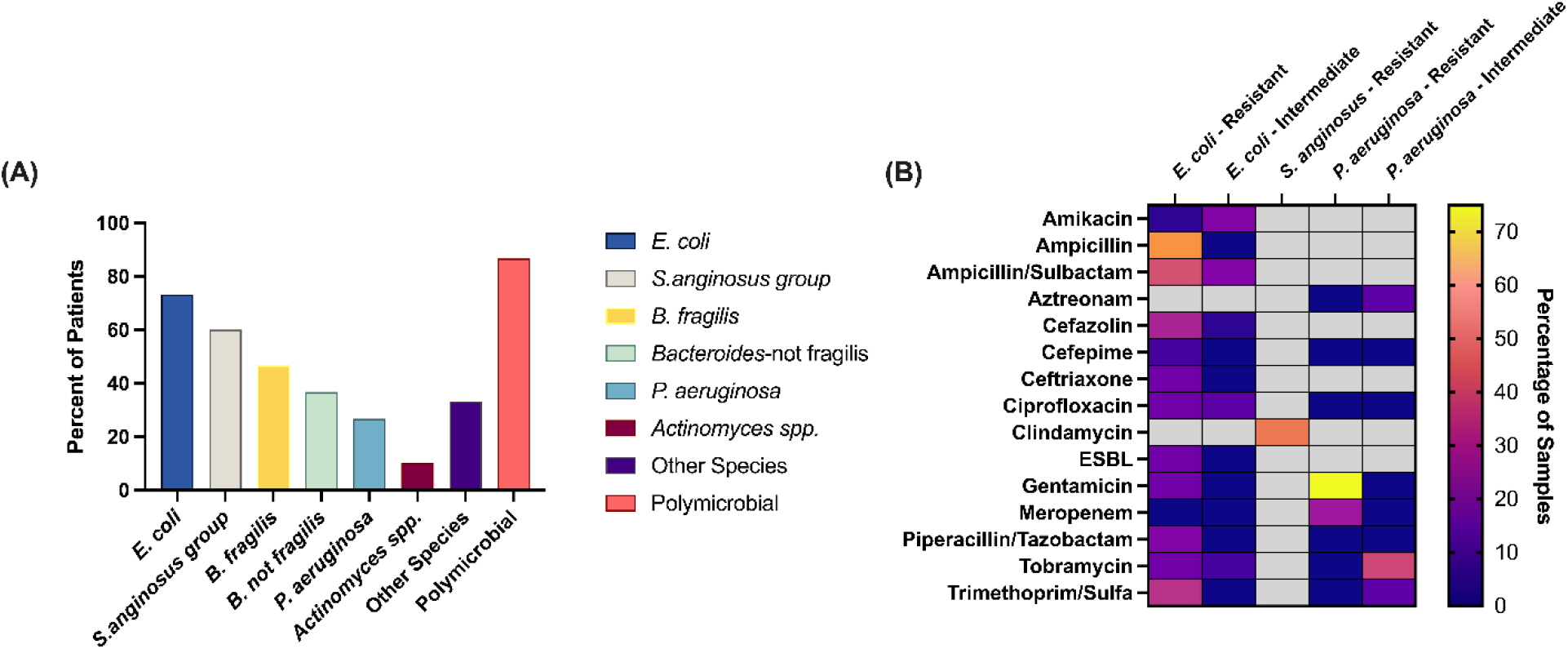
Bacterial Species in Perforated Appendicitis. (A) Percentage of patients with each bacterial species present from intraoperative cultures. (B) Heat map indicating the frequency of resistance to each antibiotic for the three most prevalent aerobic species.

### Antibiotic resistance

The clinical microbiology labs reported sensitivities in 92.5% (25/27) of *E. coli* strains, 22.2% (4/18) of *S. anginosus*, and 88.9% (8/9) of *P. aeruginosa*. Antibiotic sensitivities were unavailable from the clinical microbiology lab for 77.8% (14/18) of *S. anginosus* group strains, as *S. anginosus* is not routinely tested for antibiotic resistance at our institution.

Antibiotic resistance was common in *E. coli, S. anginosus*, and *P. aeruginosa* (Figure 2B). Antibiotic resistance was most prevalent in *E. coli* strains, with resistance to ampicillin in 56% (14/25) of strains, ampicillin-sulbactam in 40% (10/25), trimethoprim-sulfamethoxazole in 32% (8/25), cefazolin in 28% (7/25), and piperacillin/tazobactam in 20% (4/25) (Figure 2B). *E*. coli demonstrated intermediate resistance to amikacin (20%, 5/25), ampicillin-sulbactam (20%, 5/25), and ciprofloxacin (12%, 3/25). Of the 4 S. *anginosus* group strains with reported antibiotic sensitivities, 2 (50%) strains were pan-susceptible to all antibiotics tested and 2 (50%) strains demonstrated resistance to clindamycin. *Pseudomonas aeruginosa* strains were most commonly resistant to gentamicin (75%, 6/8) and meropenem (12.5%, 2/8), with intermediate resistance to tobramycin (37.5%, 3/8), aztreonam (12.5%, 1/8), and trimethoprim-sulfamethoxazole (12.5%, 1/8).

### PDT results for Gram-positive and Gram-negative bacteria

Among all 53 bacterial isolates (Figure 3A), relative to the no treatment conditions (MB−L−: 7.66±1.41 log_10_), methylene blue alone resulted in significant bacterial reduction relative to control (MB+L−: 6.37±2.17 log_10_, p<0.001, Figure 3A), whereas laser light alone did not (MB−L+: 7.45±1.49 log_10_, p=0.49). In the PDT-treated isolates, MB-PDT (MB+L+: 2.51±2.17 log_10_) resulted in an overall 5.72±0.39 log_10_ reduction in bacterial burden (p<0.001, Figure 3A), relative to the no treatment condition.

**Figure 3.**
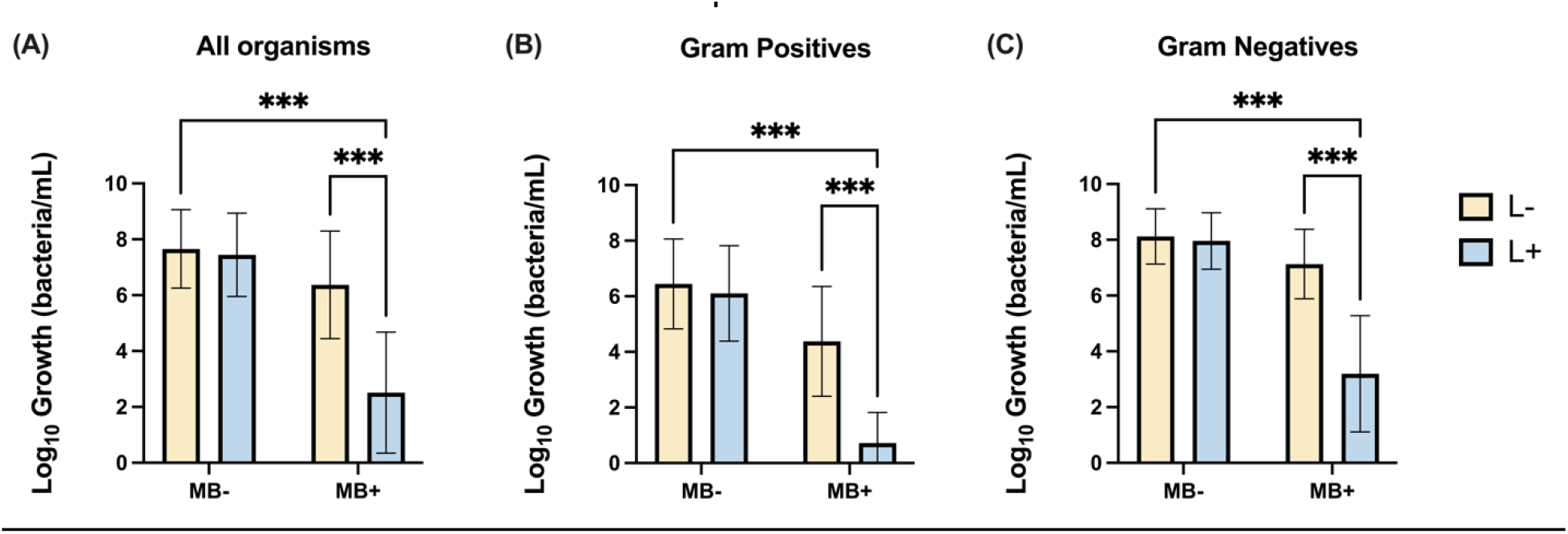
MB-PDT (MB+L+) is effective in reducing bacterial burden in (A) all organisms, (B) gram-positive, and (C) gram-negative bacteria isolated from human subjects with perforated appendicitis. MB+: treated with methylene blue; MB-: not treated with methylene blue, L+: treated with light; L−: not treated with light. Analyzed with 2-way ANOVA, *** indicates p<0.001.

Among the 15 Gram-positive isolates (Figure 3B), which included both *S. anginosus* group and *S. capitis*, relative to the no treatment condition (MB−L−: 6.45±1.62 log_10_) methylene blue without laser light resulted in a significant reduction in bacterial burden (MB+L−: 4.38±1.97 log_10_, p<0.001), whereas laser light alone did not reduce bacterial burden (MB−L+: 6.10±1.72 log_10_, p=0.74). The MB-PDT-treated (MB+L+: 0.72±1.01 log_10_) Gram-positive bacteria showed a significant bacterial burden reduction of 5.72±0.93 log_10_ (p<0.001, Figure 3B) compared to the no-treatment condition.

We noted a similar pattern among the 38 Gram-negative isolates, including both *E. coli* and *Pseudomonas aeruginosa*. Compared to no treatment (MB−L−: 8.12±0.99 log_10_), methylene blue without laser light significantly reduced bacterial burden (MB+L−: 7.13±1.25 log_10_, p<0.001), whereas laser light alone did not (MB−L+: 7.96±1.01 log_10_, p=0.75). The MB-PDT-treated Gram-negative (MB+L+: 3.20±2.09 log_10_) bacteria exhibited a substantial total reduction of 4.92±0.42 log_10_ (p<0.001, Figure 3C).

### PDT results for individual bacterial species

#### Escherichia coli

Using 27 *E. coli* isolated strains, compared to no treatment (MB−L−: 8.14±1.05 log_10_), methylene blue without laser light resulted in a significant bacterial reduction relative to control (MB+L−: 6.95±1.32 log_10_, p<0.001, Figure 4A), while laser light alone did not (MB−L+: 7.93±1.11 log_10_, p=0.49). MB-PDT (MB+L+: 2.28±1.28 log_10_) caused a substantial bacterial reduction compared to no treatment, with an overall 5.86±0.39 log_10_ reduction in bacterial burden (p<0.001, Figure 4A).

**Figure 4.**
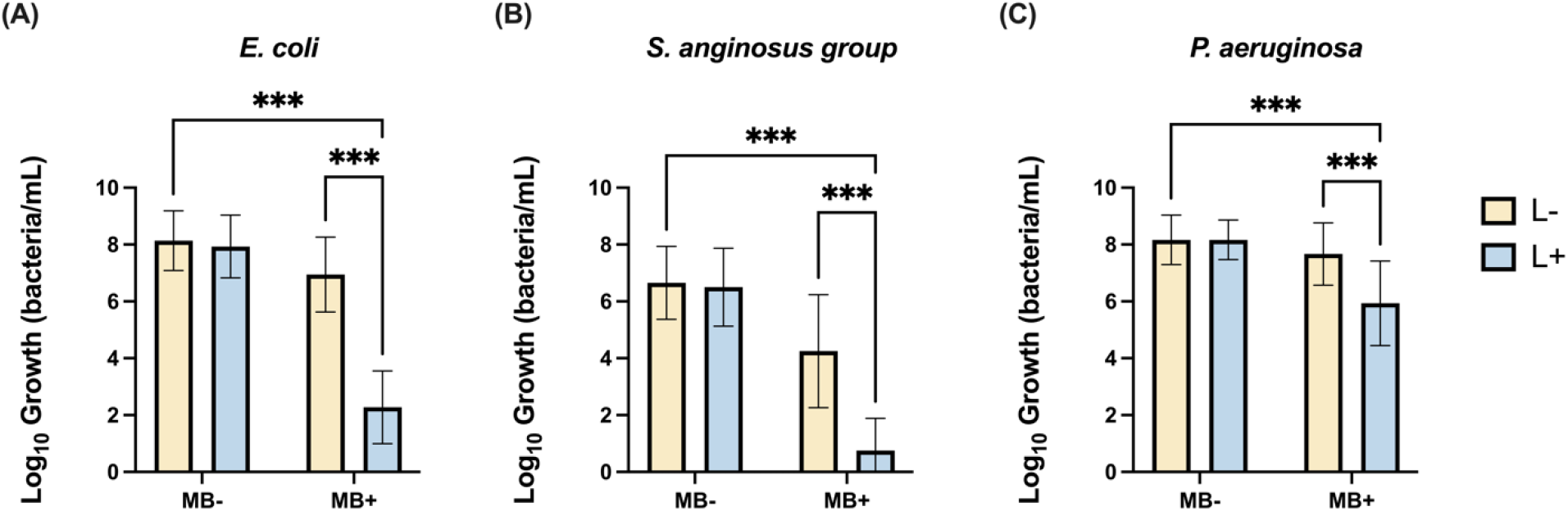
MB-PDT (MB+L+) is effective against (A) *Escherichia coli* (n = 26), (B) *Streptococcus anginosus* (n = 16), and (C) *Pseudomonas aeruginosa* (n = 9) isolates from human pediatric perforated appendicitis. MB+: treated with methylene blue; MB-: not treated with methylene blue, L+: treated with light; L−: not treated with light. Analyzed with 2-way ANOVA, *** indicates p<0.001.

#### Streptococcus anginosus group

Across 17 tested strains, compared to no treatment (MB−L−: 6.68±1.23 log_10_), the burden of *S. anginosus* was significantly reduced with methylene blue alone (MB+L−: 4.16±1.94 log_10_, p<0.001, Figure 4B), while bacterial burden for *S. anginosus* treated with laser light alone did not decrease (MB−L+: 6.51±1.37 log_10_, p=0.96). *S. anginosus* treated with MB-PDT (MB+L+: 0.65±1.09 log_10_) experienced near complete bacterial kill in all strains with minimal variation, resulting in an overall 5.91±0.88 log_10_ bacterial reduction (p<0.001, Figure 4B).

#### Pseudomonas aeruginosa

Empirical observation showed a less robust response to MB-PDT for *Pseudomonas* and increased variation among strains. Compared to no treatment (MB−L−: 8.15±0.92 log_10_), neither methylene blue alone nor laser light alone resulted in a significant bacterial reduction compared to no treatment (MB+L−: 7.58±1.14 log_10_, p=0.17; MB−L+: 8.19±0.69 log_10_, p>0.99). However, MB-PDT (MB+L+: 5.69±1.44 log_10_) treatment of *Pseudomonas aeruginosa* resulted in a significant 2.23±0.64 log_10_ bacterial reduction, compared to the no treatment condition (p<0.001, Figure 4C).

Due to the substantial variability in MB-PDT effectiveness against *Pseudomonas aeruginosa*, additional experiments were conducted using strains initially less susceptible to the standard MB-PDT obtained from two patients. Each sample was exposed to the no-treatment and control conditions, along with MB and 665 nm laser light at fluences of 7.2, 14.4, and 28.8 J/cm^2^. Using MB-PDT and increasing the fluence did not reliably increase bacterial kill for *Pseudomonas aeruginosa* (Figure 5).

**Figure 5.**
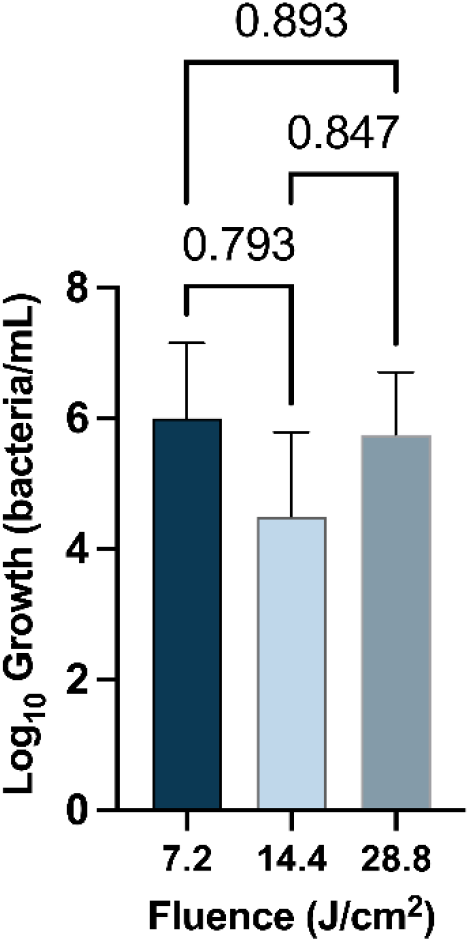
Increased laser light fluences does not significantly change MB-PDT (MB+L+) effectiveness in two *P. aeruginosa* strains from pediatric patients with perforated appendicitis. Connecting bars indicate p values for pairwise comparisons.

### Comparing PDT results for antibiotic-resistant and susceptible *E*. *coli* strains

In 20 antibiotic-resistant *E. coli* strains, compared to no treatment (M-L−: 8.18±1.13 log_10_), methylene blue without laser light resulted in a significant bacterial reduction (MB+L−: 7.13±1.02 log_10_, p<0.001), while laser light alone did not (MB−L+: 7.93±1.27 log_10_, p=0.505). MB-PDT (MB+L+: 2.48±1.12 log_10_) caused a substantial 5.70±0.67 log_10_ reduction in bacterial burden in antibiotic-resistant *E. coli* strains, compared to no treatment (p<0.001, Figure 6A).

**Figure 6.**
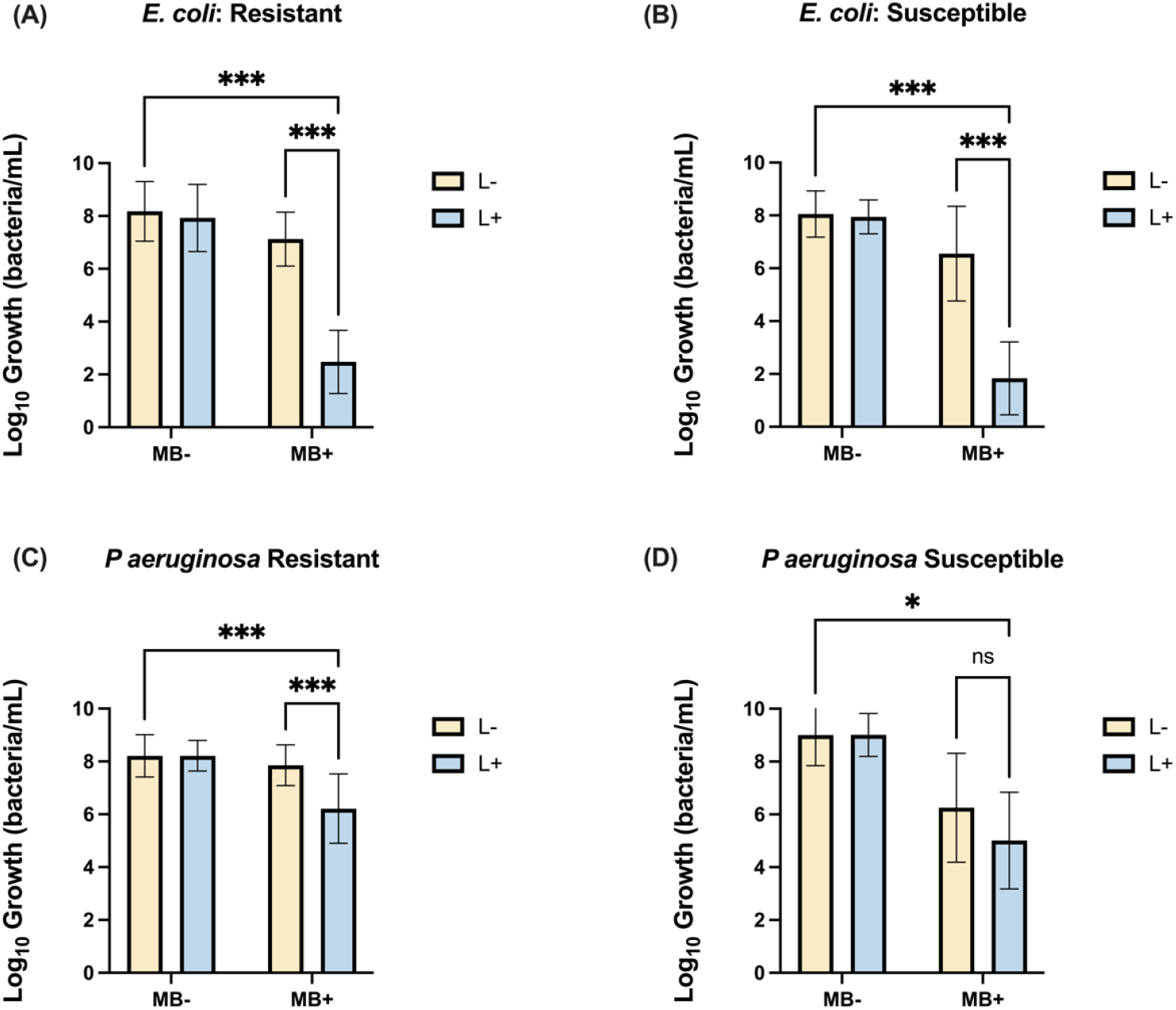
MB-PDT is effective at treating (A) *E. coli* strains with antibiotic resistance (n = 20), (B) *E. coli* strains without antibiotic resistance (n = 9), (C) *P. aeruginosa* strains with antibiotic resistance (n = 7), and (D) *P. aeruginosa* strains without antibiotic resistance (n = 2). MB+: treated with methylene blue; MB-: not treated with methylene blue, L+: treated with light; L−: not treated with light. Analyzed with 2-way ANOVA, *** indicates p<0.001; *p<0.05; ns indicates p≥0.05

In 9 antibiotic pan-susceptible *E. coli* strains, methylene blue without laser light resulted in a significant bacterial reduction (MB+L−: 6.56±1.79 log_10_, p<0.001), while laser light alone did not (MB−L+: 7.94±0.64 log_10_, p=0.960), compared to no treatment (MB−L−: 8.06±0.87 log_10_).

Treatment with MB-PDT (MB+L+: 1.83±1.38 log_10_) caused a substantial overall 6.22±1.21 log_10_ reduction in antibiotic pan-susceptible strains compared to no treatment (p<0.001, Figure 6B).

When we compared the MB-PDT response of antibiotic-resistant (MB+L+, R) to pan-susceptible (MB+L+, NR), *E. coli* strains, we did not observe a significant difference in the magnitude of bacterial kill (p=0.550, Supplemental Figure S2A).

### Comparing PDT results for antibiotic-resistant and susceptible *S*. *anginosus* strains

Compared to no treatment (MB−L−: 5.50±1.00 log_10_), in 2 antibiotic-resistant *S. anginosus* strains, the bacterial burden was significantly reduced with methylene blue alone (MB+L−: 2.00±0.82 log_10_, p<0.05). *S. anginosus* isolates treated with laser light alone did not decrease bacterial burden (MB−L+: 4.50±0.58 log_10_, p=0.245). MB-PDT-treated strains (MB+L+: 1.00±1.16 log_10_) demonstrated an overall 4.50±4.62 log_10_ reduction in bacterial burden that did not reach statistical significance (p=0.054). Antibiotic-resistant (R) *S. anginosus* strains did not have a significantly different response to treatment with MB-PDT (MB+L+) compared to antibiotic-susceptible (NR) strains (p>0.999, Supplemental Figure S2B).

### Comparing PDT results for antibiotic-resistant and susceptible *P*. *aeruginosa* strains

In 7 antibiotic-resistant *P. aeruginosa* strains, compared to no treatment (MB−L−: 8.21±0.80 log_10_), neither methylene blue alone nor laser light alone resulted in a significant bacterial reduction (MB+L−: 7.86±0.77 log_10_, p=0.307; MB−L+: 8.21±0.58 log_10_, p>0.999). However, treatment with MB-PDT (MB+L+: 6.21±1.31 log_10_) resulted in a substantial 2.00±1.192 log_10_ bacterial reduction compared to the no treatment control condition (p<0.01, Figure 6C).

In 2 antibiotic pan-susceptible *P. aeruginosa* strains, compared to no treatment (MB−L−: 9.00±1.16 log_10_), there was no difference in bacterial reduction in groups treated with methylene blue alone, laser light alone, nor MB-PDT (MB+L−: 6.25±2.06 log_10_, p=0.300; MB−L+: 9.00±0.82 log_10_, p>0.999; MB+L+: 9.21±5.21 log_10_, p=0.098, Figure 6D). Furthermore, there was no difference in response to treatment with MB-PDT (MB+L+) in antibiotic-resistant (R) *P. aeruginosa* strains compared to those pan-susceptible (NR) to the tested antibiotics (p=0.473, Supplemental Figure S2C).

## Discussion

This is the first study demonstrating the feasibility and effectiveness of PDT against the most common microbes isolated from intraoperative aspirates in pediatric patients with perforated appendicitis. Despite frequent antibiotic resistance in bacterial strains, treatment with MB-PDT resulted in >99.9% cell kill for *E. coli* and *Streptococcus anginosus* group. While MB-PDT was effective against *Pseudomonas aeruginosa*, the treatment effect was less robust, and more variable compared to *E. coli* and *S. anginosus group*. As rates of antibiotic resistance increase^33^, MB-PDT offers the potential of a non-antibiotic solution to treating the contaminated abdomen for infectious source control.

Multiple studies have shown that perforated appendicitis often results in a polymicrobial infection, with *E. coli, Bacteroides fragilis, Streptococcus* species, and *P. aeruginosa* being the most common organisms isolated from perforated appendicitis^35–39^. We demonstrated similar bacterial prevalence among patient cultures, with *E. coli, S. anginosus group, Bacteroides fragilis*, and *P. aeruginosa* being the most common bacteria in our samples, and most patients having polymicrobial cultures (86.7%).

Other studies have investigated the antimicrobial efficacy of PDT, including utilization of the photosensitizer methylene blue^12^. These studies have demonstrated the effectiveness of PDT in preclinical and clinical settings for other indications such as periodontal disease and chronic ulcers^34–36^. Baran *et al*. have previously demonstrated the safety and efficacy of treatment with MB-PDT in established abscess cavities^37,38^. These studies have shown that antibiotic-resistant strains are vulnerable to PDT^39–41^. However, these applications have often been limited in scope, focusing on *in vitro* treatment of individual bacterial species^42^. Further, most prior literature has utilized high fluence and fluence rates, even for the treatment of planktonic culture. For example, Piksa *et al*. described modes of 100 mW/cm^2^ and 30 J/cm^2^ across a wide range of reported values in the literature^43^. Here, we demonstrated high efficacy across a broad spectrum of bacterial species at a much lower fluence rate (4 mW/cm^2^) and fluence (7.2 J/cm^2^).

Exploration of these lower light dose parameters is critical for eventual clinical application, as delivery of high fluence rates to an extended surface, such as the peritoneal cavity, can be challenging. In simulation studies, we have shown that lower fluence rates increase patient eligibility for PDT^44,45^.

We found that MB-PDT resulted in a >99.9% antimicrobial effect against Gram-positive and Gram-negative bacteria. While there was variability among bacterial species, the 3 log_10_ reduction threshold was surpassed for all bacteria tested except *P. aeruginosa*. For the most prevalent species, *E. coli* and *S. anginosus* group, treatment with MB-PDT resulted in a decrease of greater than 5.3 and 5.0 log_10_, respectively.

There were substantial empirical differences in *P. aeruginosa’s* response to MB-PDT. The mean reduction was below the hypothesized threshold; however, the observed 2.46 log_10_ reduction was within the 95% confidence interval. The variability in response to MB-PDT between *P. aeruginosa* strains led us to test these bacteria with different light fluences. Hawryluk *et al*. demonstrated increased MB-PDT efficacy against *P. aeruginosa* when fluence was increased over a comparable range to the one studied here^31^. However, we demonstrated no difference in bacterial reduction in the two *P. aeruginosa* strains treated with MB-PDT at 7.2 J/cm^2^, 14.4 J/cm^2^, and 28.8 J/cm^2^ fluence. *P. aeruginosa* shows considerable variability between strains regarding metabolic activity, antibiotic resistance profile, and secretion of extracellular pigments^46^. We observed various phenotypes in our *P. aeruginosa* strains, including pyocyanin and pyoverdine producers. Therefore, differences in MB-PDT efficacy may be strain-specific, highlighting the importance of setting universally efficacious PDT dose parameters.

There is controversy in the literature regarding the contribution of antibiotic-resistant microbes to the development of postoperative intrabdominal abscesses. Gonzales *et al*. demonstrated no association between antibiotic resistance and post-operative abscess development^36^, whereas others have reported data from intrabdominal abscess cultures with antibiotic-resistant pathogens, including *Pseudomonas aeruginosa* and *E. coli*^37,38^. Since the incidence of multidrug-resistant strains continues to rise, we specifically evaluated the efficacy of MB-PDT against antibiotic-resistant strains. We found that antibiotic-resistant *E. coli, S. anginosus group*, and *P. aeruginosa* strains had similar bacterial reduction from MB-PDT as antibiotic-susceptible strains.

In a distinct but related patient population, our group performed a Phase 1 clinical trial using MB-PDT to treat abdominopelvic abscesses in adults^37,38^. This clinical trial demonstrated a shorter duration of both clinical symptoms and fever in subjects who received higher PDT light doses, with no study-related adverse or serious adverse events observed. These encouraging clinical results highlight the translational potential of MB-PDT for treating intra-abdominal infection, but are limited to well-defined abscess cavities. With these data and the data from the current study, we envision that MB-PDT could be used to treat the entire peritoneal cavity during an operation associated with bacterial contamination (such as an appendectomy or colon resection for diverticulitis) to reduce or eliminate the bacterial population present in the peritoneal cavity. This would involve the local delivery of MB followed by targeted illumination using fiber optic delivery. As most appendectomies are performed laparoscopically, these tasks could be achieved through standard laparoscopic trocars. MB-PDT would, therefore, require nominal additional time and equipment in the operating room while potentially reducing complications and hospitalization duration relative to the current standard of care. A similar approach, albeit with an open surgical technique, has been employed for intra-operative PDT treatment of mesothelioma^47^.

We acknowledge some limitations in the current study. To allow comparison between bacterial strains and to previous results^30^, only a single combination of drug and light dose was explored for most bacterial samples. Higher light fluence has been shown to improve PDT response^31^, particularly for Gram-negative bacteria. Optimization of treatment parameters will, therefore, be explored in future studies. Additionally, we only examined planktonic and monomicrobial cultures. Like Gram-negative bacteria grown planktonically, bacterial biofilms may require increased drug or light doses to achieve comparable PDT efficacy to the results reported here. As most of the patients in this study had polymicrobial infections, further examination of PDT efficacy in polymicrobial cultures will be an essential step toward clinical translation. In addition, anaerobic bacteria were not submitted to *in vitro* PDT experiments. Successful PDT requires the presence of molecular oxygen for conversion into cytotoxic reactive oxygen species. These levels of molecular oxygen would be inherently toxic to anaerobic bacteria, so these species were excluded from the current analysis.

In conclusion, we have demonstrated that MB-PDT at low light dose is effective against bacteria isolated from pediatric patients with perforated appendicitis. These results motivate further investigation of MB-PDT in the context of perforated appendicitis. Based on these *in vitro* results and our clinical experience with MB-PDT^38^, we are optimistic that this treatment could substantially reduce the intra-abdominal infection secondary to PA and, ultimately, reduce associated complications and hospital length of stay for patients with PA. This proof-of-concept study establishes a strong foundation to motivate MB-PDT as a novel adjunct to surgery, potentially reducing hospital stays for parenteral antibiotics and the development of postoperative abscesses.

## Acknowledgments

The authors would like to thank Dr. Laurel Baglia for their assistance in preparing photodynamic therapy protocols. This work was funded by grant R21 AI178152 from the National Institutes of Health and the National Institute of Allergy and Infectious Diseases.

## Data Availability

The datasets generated and analyzed during the current study are available from the corresponding author upon reasonable request.

