## Supplemental Figures for "Methylene Blue-Photodynamic Therapy Effectively Kills Antibiotic-Resistant Bacteria from Pediatric Patients with Perforated Appendicitis"

**Figure S1. Two-way ANOVA demonstrating efficacy of MB-PDT *Staphylococcus capitis* isolate.** MB+: treated with methylene blue; MB-: not treated with methylene blue, L+: treated with light; L-: not treated with light. Analyzed with 2-way ANOVA, ** indicates $p<0.002$. Using a single isolate that was antibiotic-susceptible, compared to no treatment (MB-L-: 4.83±2.14 $\log_{10}$), neither methylene blue alone nor laser light alone resulted in a significant bacterial reduction (MB+L-: 4.500±1.87 $\log_{10}$, p=0.935; MB-L+: 4.33±2.16 $\log_{10}$, p=0.822). However, MB-PDT (MB+L+: 0.00±0.00 $\log_{10}$) resulted in near complete bacterial kill (MB+L+: 4.83±2.13 $\log_{10}$, $p<0.002$).

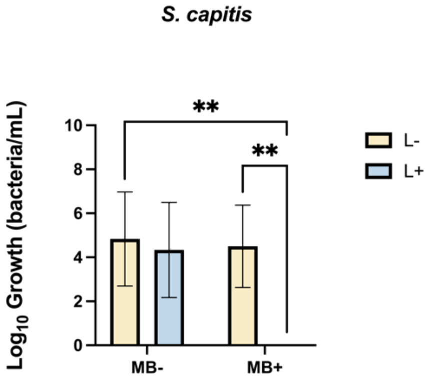


**Figure S2. There is no significant difference in MB-PDT response in antibiotic-susceptible versus antibiotic-resistant (A) *E. coli* (p = 0.760), (B) *S. anginosus* (p > 0.999), and (C) *P. aeruginosa* (p = 0.991).** Analyzed with 1-way ANOVA.

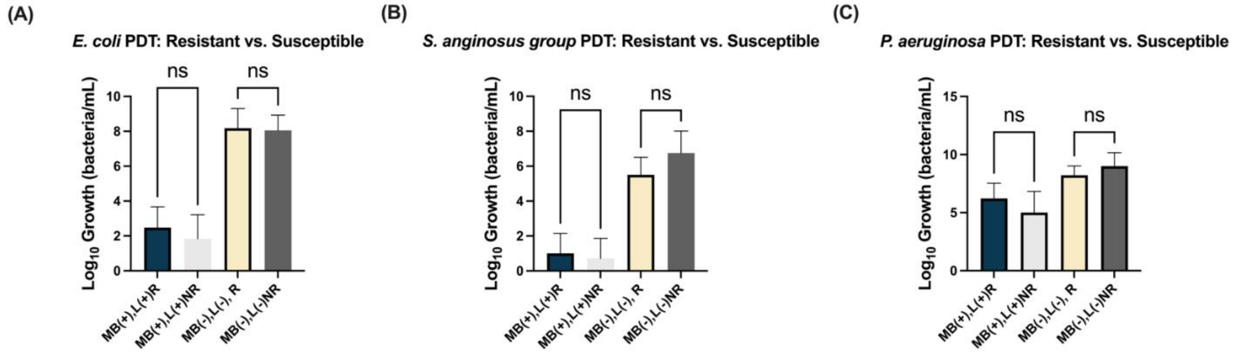
